# Is biomedical research self-correcting? Modeling insights on the persistence of spurious science

**DOI:** 10.1101/2023.07.17.549436

**Authors:** David Robert Grimes

## Abstract

The reality that volumes of published research are not reproducible has been increasingly recognised in recent years, notably in biomedical science. In many fields, spurious results are common, reducing trustworthiness of reported results. While this increases research waste, a common response is that science is ultimately self-correcting, and trustworthy science will eventually triumph. While this is likely true from a philosophy of science perspective, it does not yield information on how much effort is required to nullify suspect findings, nor factors that shape how quickly science may be correcting in the publish-or-perish environment scientists operate. There is also a paucity of information on how perverse incentives of the publishing ecosystem, which reward novel positive findings over null results, shaping the ability of published science to self-correct. Precisely what factors shape self-correction of science remain obscure, limiting our ability to mitigate harms. This modelling study illuminates these questions, introducing a simple model to capture dynamics of the publication ecosystem, exploring factors influencing research waste, trustworthiness, corrective effort, and time to correction. Results from this work indicate that research waste and corrective effort are highly dependent on field-specific false positive rates and the time delay before corrective results to spurious findings are propagated. The model also suggests conditions under which biomedical science is self-correcting, and those under which publication of correctives alone cannot stem the propagation of untrustworthy results. Finally, this work models a variety of potential mitigation strategies, including researcher and publication driven interventions.

**Significance statement:** In biomedical science, there is increasing recognition that many results fail to replicate, impeding both scientific advances and trust in science. While science is self-correcting over long time-scales, there has been little work done on the factors that shape time to correction, the scale of corrective efforts, and the research waste generated in these endeavours. Similarly, there has been little work done on quantifying factors that might reduce negative impacts of spurious science. This work takes a modeling approach to illuminate these questions, uncovering new strategies for mitigating the impact of untrustworthy research.

## Introduction

Biomedical science is integral to human prosperity. Diligent research keeps us healthy, functioning as a compass towards a better future for humanity. Yet despite its critical importance to our collective well-being, there are serious issues with how biological science and medicine are currently practiced. This was illustrated starkly by the on-going COVID-19 pandemic, which saw an explosion of COVID-related research. This uptick in publication not only highlighted how science is practiced, but raised concerns over its failings. In a short period, much COVID research subsequently shown to be of low methodological quality, and unsuitable for drawing inferences^1–6^. Some of these failings were high-profile enough to alter public opinion and even endanger public health. In 2020, Donald Trump proclaimed hydroxychloroquine a panacea for COVID-19 based on flawed analysis, and later works extolling anti-parasitic medicine Ivermectin for COVID ignited world attention, persisting despite later analysis exposing them as unreliable. Consequently, millions worldwide embraced ineffectual medicines with deleterious consequences, spending hundreds of millions of dollars on an unsuitable treatments with a potential of harm^7,8^.

Irreproducible research was already a problem long before the dawn of COVID. Swathes of ostensibly important biomedical results simply do not withstand deeper scrutiny^9^. In fields as diverse as psychology to genomics to cancer research, there has been increasing recognition that we are experiencing a pan-field replication crisis, where seemingly important results do not hold under inspection. This is an especially pronounced problem in medicine, when life-or-death clinical decisions might pivot on research outcomes. There are many culprits for this state of affairs. Inappropriate statistical practices and manipulation underpin many spurious results^10–14^, affecting up to 75% of biomedical publications^15–17^. In fields as vital as cancer research, only 11% − 50% of preclinical results are deemed replicable^18,19^. This is compounded by publication bias, where high-prestige journals fixate upon novel “positive” results over more diligent null results. The metric-driven “Publish-or-Perish” environment in which scientists and physicians inevitably operate is also a factor, as this inadvertently privileges dubious findings^20^, dubbed “*the natural selection of bad science*”^21^. Modeling studies also suggest that scientists acting rationally to maximize career value of their publications leads to a high rate of false positives^22^ Data and methods are often jealously guarded, making it difficult to spot methodological errors or even verify assertions made in published literature. In many instances, even seemingly robust results transpire to be fragile and not robust^23,24^.

Publication bias also means that for many scientists, there is little reward to be had in pursing corrective endeavours, and only minor benefit in disseminating null-results. Consequently, replication efforts are rarely undertaken and null results are often buried and not shared for publication. This “file-drawer” problems means that important and diligent negative findings are often not submitted for publication, whilst spurious findings are unduly favoured^25–28^. This is on the face of it a strange state of affairs - it is every bit as critical to know a particular treatment had no effect as to know it has one, but only the latter is highly valued. While modelling suggests this situation inherently privileges careless or unethical researchers over more diligent ones, a common response is that science is ultimately self-correcting, and thus such errors will eventually be remedied. Self-correction itself lies at the heart of philosophy of science, considered a vital hallmark of scientific practice. Science by definition pivots on empirical observations, and holds its results to perpetual scrutiny. With the passage of time, a spurious result will fail replication, and the scientific record will be corrected.

This is likely true on a long enough timescale, but it does not take into account the substantial obstacles impeding correction. Science is an enterprise conducted by humans, with all of their incentives and failing. If positive results are disproportionately rewarded while replication and null findings systematically impeded, than vital corrective efforts will be stunted. In recent years, many authors have questioned whether our current paradigm truly enables self-correction. This has been particularly examined in the context of psychology, one of the first major fields to identify a replication crisis and examine its impacts objectively^29,30^. In other fields, there is marked reticence from journals to correct flawed works^31^. One recent review noted that biomedicine as a field was severely lacking in appropriate levels of transparency and, crucially, critical appraisal. The authors conclusion was that fields without transparency and and robust, verifiable mechanisms for critical appraisal “*cannot reasonably be said to be self-correcting, and thus do not warrant the credibility often imputed to science as a whole*”^32^.

Accordingly, science as it currently operates is not always self-correcting, causing harm to both science and society^33^. Scientific practice is also frequently to blame. Examining the history of medical research, Doug Altman and Iveta Simera detailed how poor experimental design and subpar reporting was a longstanding problem, writing that “*the quality of published papers is a fair reflection of the deficiencies of what is still the common type of clinical evidence. A little thought suffices to show that the greater part cannot be taken as serious evidence at all*.”^34^. Despite high profile expert guidelines on research conduct, many published papers still demonstrate substandard reporting and methodological issues^35,36^. Untrustworthy findings that do not withstand closer inspection are ultimately research waste, contributing to noise without benefiting patients or the research community^37^. By some estimates, research waste is considered to constitute up to 85% of biomedical science output^38^. Even when retracted, these now-phantom publications can remain highly cited in literature long after their credibility has decayed^39–45^.

While the problem of self-correction has begun to garner significant attention, there is as yet no means to quantify the likely corrective effort required to sustain self-correction, nor the volume of research waste likely to be produced in a given biomedical field. There is no literature to the author’s knowledge that estimates the likely time taken to correction, nor the situations where self-correction may breakdown. This is not surprising, as the publishing ecosystem in which science operates is complex and multi-faceted, and direct measurement of relevant factors is difficult or impossible. Without this knowledge, however, we cannot realistically estimate the extent of problems, nor even optimally gauge potential strategies to mitigate the problem. Modeling provides model insight for such problems, and previous investigations have investigated how pressures on scientists can lead to spurious findings^20–22^ but there have been few direct investigations into how the scientific publishing ecosystem itself might modulate this problem, and what factors might ameliorate or exacerbate research waste. In this work, we take a novel mathematical modelling approach, crafting a simple dynamic model of scientific publishing and self-correction. The model derived allows to estimate the likely impacts of different factors on scientific trustworthiness, research waste, correction time, and corrective effort, subject to reasonable field-specific assumptions. The net advantage is that this not only allows us to make quantified predictions about the dynamics of scientific research, it also yields insight into possible strategies to mitigate spurious research, the ramifications of which are developed and discussed.

## Methods

### Model outline

To investigate factors shaping the research ecosystem, we consider a simple ‘gold rush’ model, where spurious results, *x*(*t*), beget consequent spurious results as other groups or the initial investigators pursue similar avenues. This increases the volume of untrustworthy literature, but eventually spurns other investigators to examine these claims and publish contrary literature from more diligent experimentation, *y*(*t*). The growth rate of consequent spurious claims is *g*. The rate at which corrective literature emerges based on the prevalence of suspect literature is *e*, and corrective attempts have a relative impact of *d*. These constants subsume different quantities, and will be defined shortly in context. The relationship between suspect and corrective literature is thus

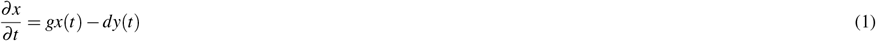

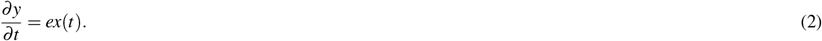

This coupled first-order differential equation system can be explicitly solved, subject to the initial condition *x*(0) = *x*_*o*_. Corrective efforts emerge after a time *t*_*d*_ from the initial erroneous publication, so that *y*(*t ≤ t*_*d*_) = 0, to account for the reality that there can be latent periods before spurious claims are challenged. This system can be solved to yield closed-formed analytical solutions for *x*(*t*) and *y*(*t*) of

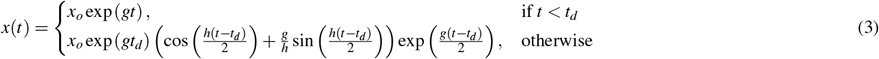

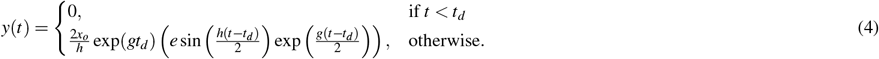

where 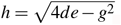. We further implement realistic parameters for *g, d*, and *e*. For a field with average research rate of *p*_*r*_ per unit time, the rate of spurious positive results depends upon the field’s false positive rate *f*_*p*_ and the publication bias of a family of relevant journals, *B*, or the fraction of significant findings published (relative to null findings) in field-specific journals. Assuming all positive results are submitted for publication, then *g* = *p*_*r*_*B f*_*p*_. Conversely, true null findings occur at a rate *e* is 1 − *f*_*p*_, with publication bias 1 − *B* for null findings. We account for the file drawer problem where only a fraction of null results are submitted for publication by introducing the submitted fraction, *s*_*r*_. It follows that *e* = *p*_*r*_(1 − *B*)(1 − *f*_*p*_)*s*_*r*_. Finally, we consider how impactful null results are relative to positive findings, given by *d* = *kp*_*r*_, where *k* is a constant value. When *k* = 1, corrective publication have equal impact to the publication ecosystem as spurious findings. Evidence to date suggests that in general, *k <* 1. When 4*de < g*^2^, corrective literature will eventually nullify spurious findings when *x*(*t*_*c*_) = 0, yielding

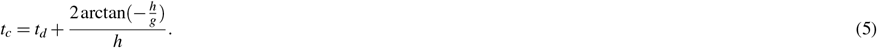

When *g*^2^ ≥ 4*de*, there is no positive real number solution for *t*_*c*_. This corresponds to the scenario where corrective literature cannot remedy the propagation of a spurious finding - a situation where science by conventional publication schema is not self-correcting.Defining research waste as the total spurious publications produced in the interval to *t*, or 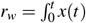 and corrective effort to nullify spurious findings as 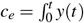, yields

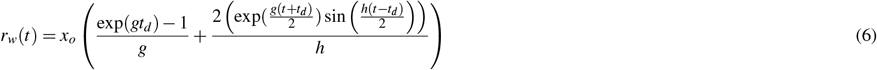

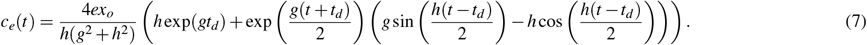

Solving these identities at *t* = *t*_*c*_ (when defined) yields the total spurious publications and corrective effort required to correct a spurious finding. For brevity, we designate *r*_*w*_(*t*_*c*_) as *r*_*w*_ and *c*_*e*_(*t*_*c*_) as *c*_*e*_ unless otherwise specified.

### Simulated scenarios

With the model established, we simulate the following scenarios:

1. **Impacts of field-specific false positive rates and corrective time-delay on result trustworthiness, correction time, and research waste. (Publishing ecosystem)**: To investigate this, we simulate the dynamics of fields with varying false positive rates (*f*_*p*_) and differing times for delaying corrective action, *t*_*d*_.
2. **Impacts of file-drawer problems on self-correction and corrective impact on research waste. (Researcher driven interventions):** To examine impacts researchers can have on the levels of research waste in a field, we can simulate the file-drawer problem by investigating effects of varying *s*_*r*_, the proportion of null results submitted. Similarly, we can also investigate the impacts of the value researchers place on null results relative to positive results by varying *k*, the relative corrective effect of null results.
3. **Hypothetical impacts of eliminating publication bias on research waste and result trustworthiness. (Publishing industry interventions):** Finally, publishers themselves have substantial scope to influence the research trustworthiness of the field through their editorial policy. In this simulation, we examine the impact of removing publication bias. In this case, there is no incentive to submit positive results. Accordingly in this hypothetical scenario, *s*_*r*_ = *k* = 1, *g* = *p*_*r*_ *f*_*p*_, and *e* = *p*_*r*_(1 − *f*_*p*_). This can be simulated for varying values of false positives *f*_*p*_ and time to corrective action, *t*_*d*_ to gauge the scope of such policy.

Sample code for these simulations is provided in electronic supplementary material.

## Results

### Impacts of field-specific false positive rates and time-delay on result trustworthiness

Figure 1 depicts the impact on the publication ecosystem of an initial spurious (*x*_*o*_ = 1) result in fields with low (*p*_*f*_ = 0.05) and high (*p*_*f*_ = 0.20) false positive rates, with simulation parameters of *p*_*r*_ = 3 submissions a year, *s*_*r*_ = 0.5 (half of all null results submitted), *B* = 0.95, and *k* = 0.6. Results are given in table 1. As expected from equation 5, *t*_*c*_ increases with false positive rate and research waste is highly dependent on time delay until corrective action, *t*_*d*_. Figure 2 shows contour plots for both corrective time *t*_*c*_ and the volume of research waste produced, *r*_*w*_. In all cases, *r*_*w*_ *> c*_*e*_ so it always takes a lesser volume of corrective work to nullify spurious results, even when *k <* 1. When *g*^2^ *>* 4*de*, there is no real value for *t*_*c*_, corresponding to a situation where spurious work cannot be remedied by corrective publication. This would occur in this example when *f*_*p*_ = 0.25, where spurious publications are irreversible by correction, illustrated in figure 3.

**Table 1.**
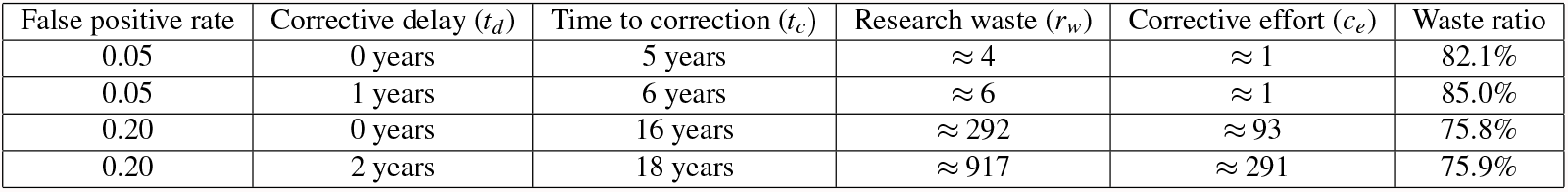
Corrective time, research waste, and corrective effort for scenarios in figure 1.

**Table 2.** Corrective time, research waste, and corrective effort for scenarios in figure 5.

| False positive rate | Corrective delay ($t_d$) | Time to correction ($t_c$) | Research waste ($r_w$) | Corrective effort ($c_e$) | Waste ratio |
| --- | --- | --- | --- | --- | --- |
| 0.05 | 0 years | 0.5 years | $\approx 1$ | $\approx 1$ | 50.4% |
| 0.05 | 1 years | 1.5 years | $\approx 1.5$ | $\approx 1$ | 78.5% |
| 0.50 | 0 years | 1 years | $\approx 1$ | $\approx 1$ | 54.3% |
| 0.50 | 2 years | 3 years | $\approx 32$ | $\approx 17$ | 66.2% |

**Figure 1.**
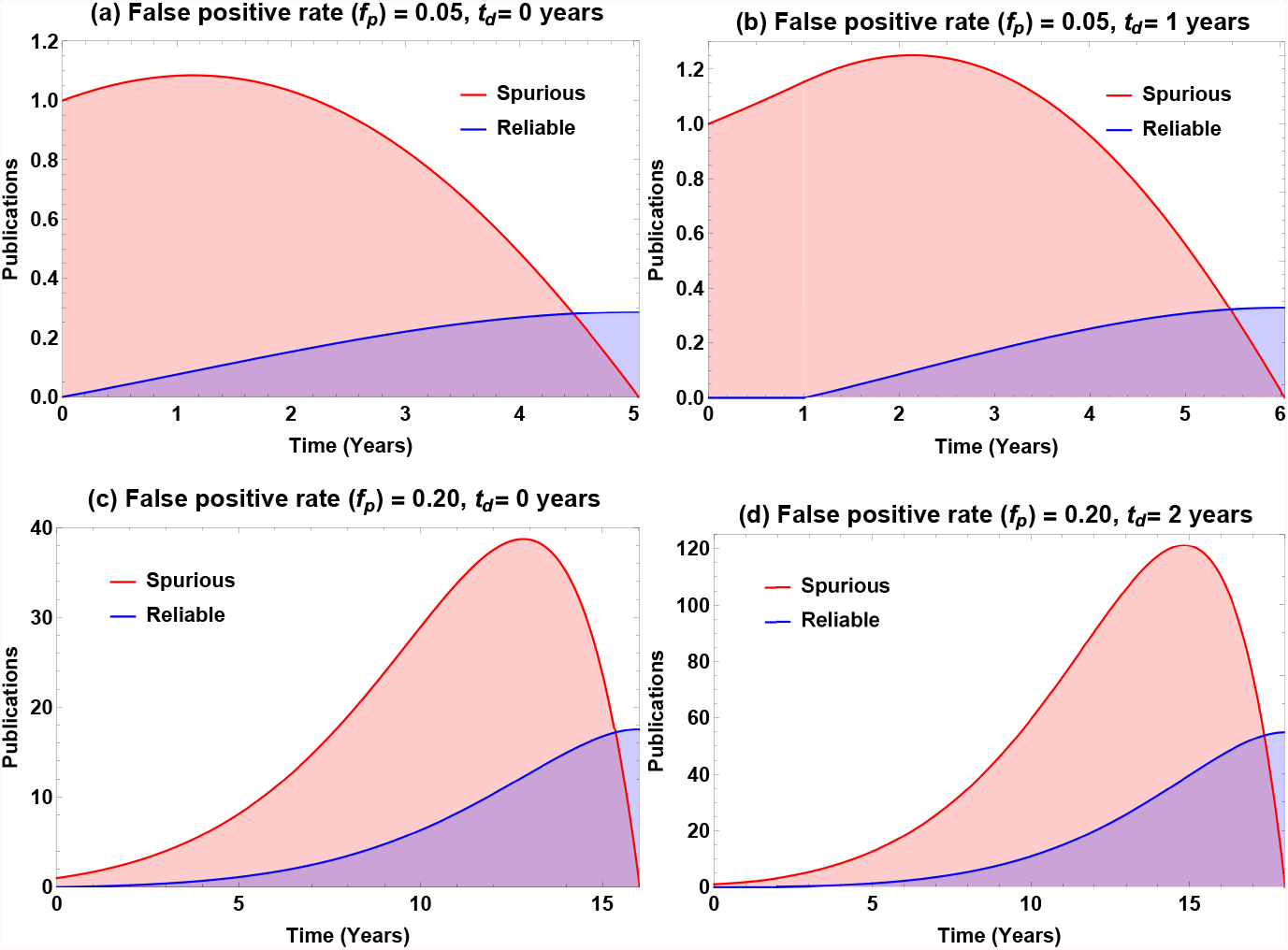
Dynamics of publishing ecosystem for (a) low false positive rate, no corrective delay (b) low false positive rate, minor corrective delay, (c) high false positive rate, no corrective delay (d) high false positive rate, 2 year corrective delay. Note the varying y-axis scales (denoting number of spurious publications released at a specific time) in all subfigures.

**Figure 2.**
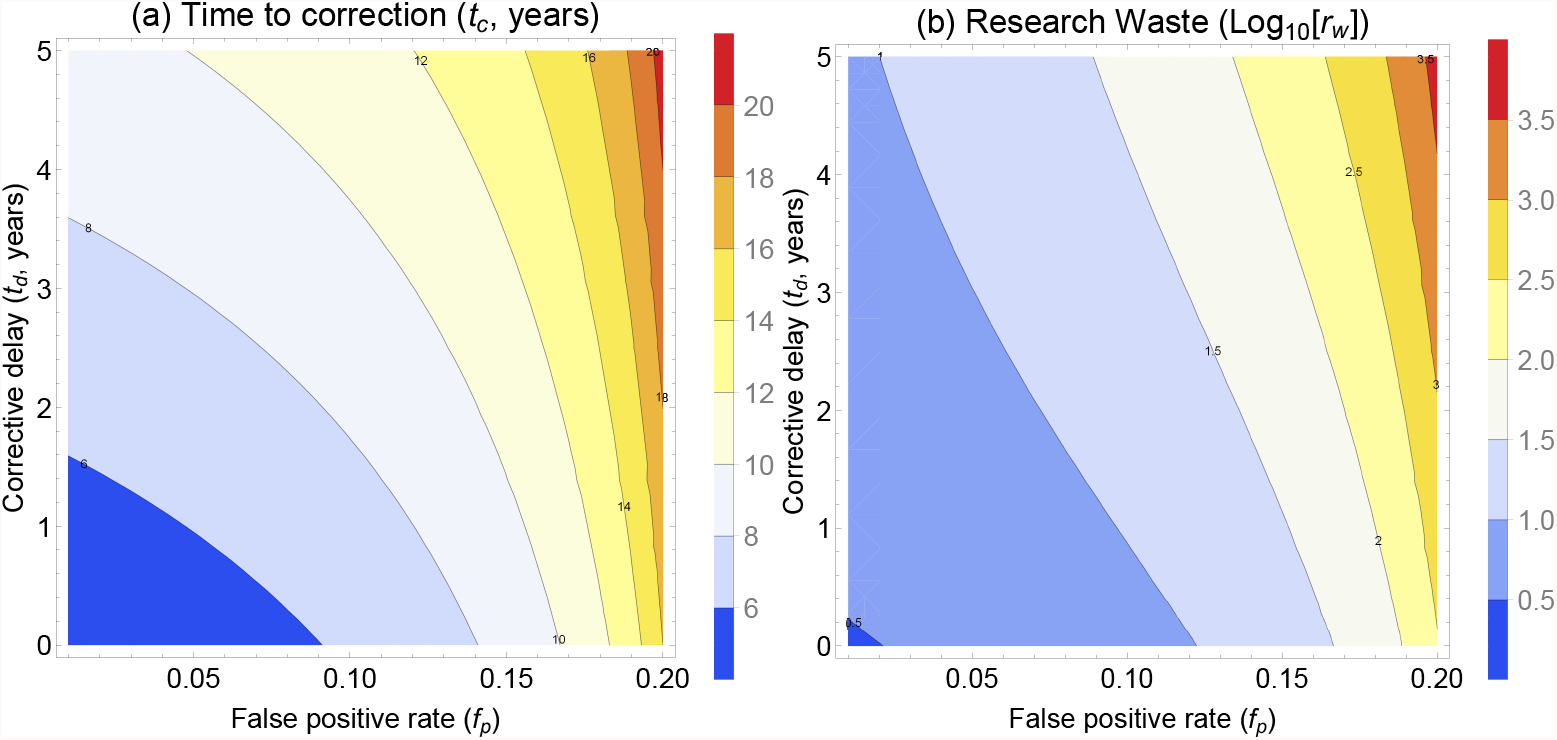
(a) Time to correction for varying values of *f*_*p*_ and *t*_*d*_ with all other parameters kept constant (b) The log 10 of research waste (*r*_*w*_) with varying values of *f*_*p*_ and *t*_*d*_ with all other parameters kept constant. Note the log scale on this figure, so that the contour line of 2 is equivalent to *r*_*w*_ = 100, 3 corresponds to *r*_*w*_ = 1000 etc.

**Figure 3.**
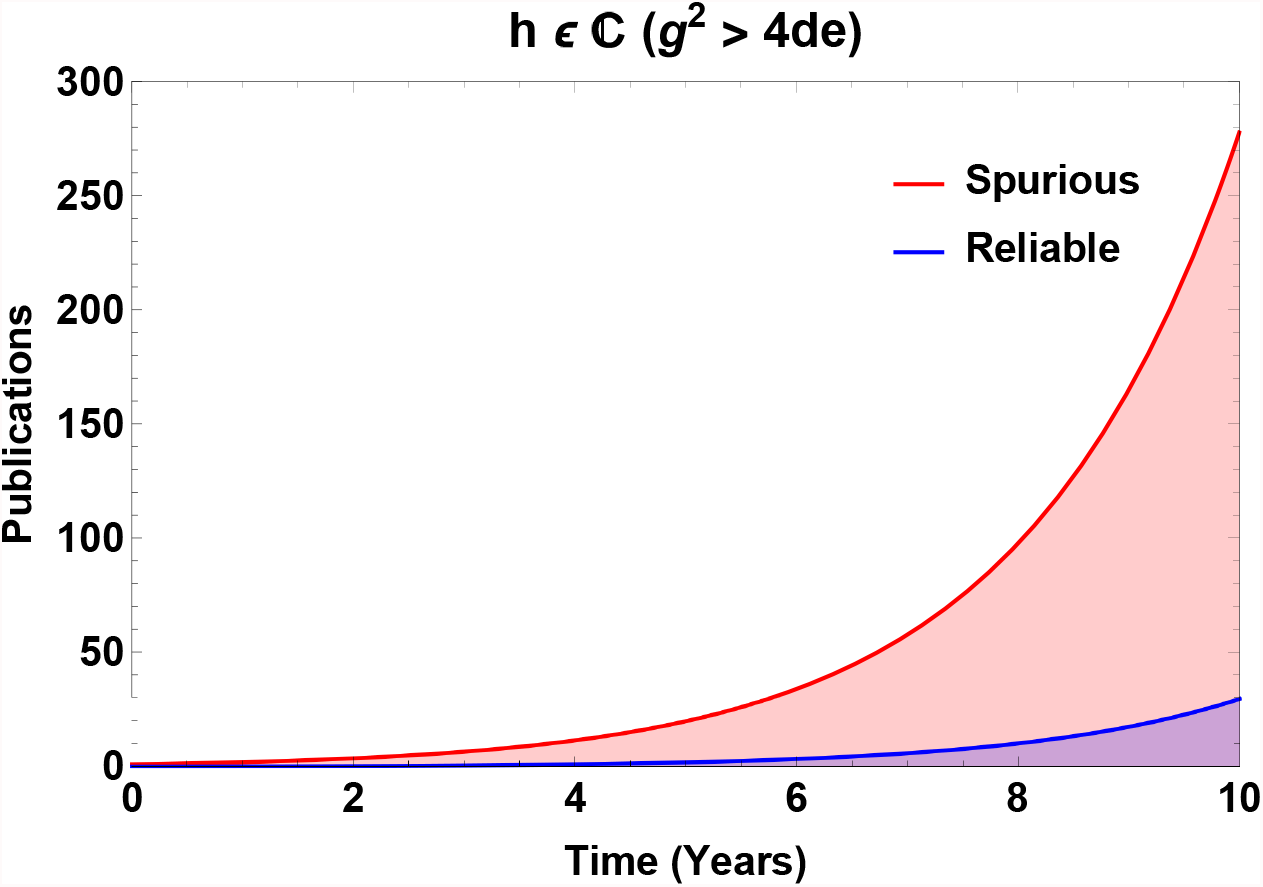
When *g*^2^ *>* 4*de*, corrective efforts cannot nullify spurious results, which grow without meaningful impediment.

### Impacts of file-drawer problems on self-correction and corrective impact on research waste

Figure 4 depicts the impact of researcher driven interventions on the time taken for correction of spurious results and the degree of research waste generated. The higher the submitted proportion of negatives (*s*_*r*_ → 1), the shorter the time to correction and less research waste generated. Similar patterns are observed when null and negative results are equally valued to positive or significant results by the research community (*k* → 1). Low values of *k* and *s*_*r*_ lead to greater volumes of research waste, more corrective effort, and longer time to correction. For sufficiently low values of *s*_*r*_ and *k, g*^2^ ≥ 4*de* and consequently no correction via publication alone is possible.

**Figure 4.**
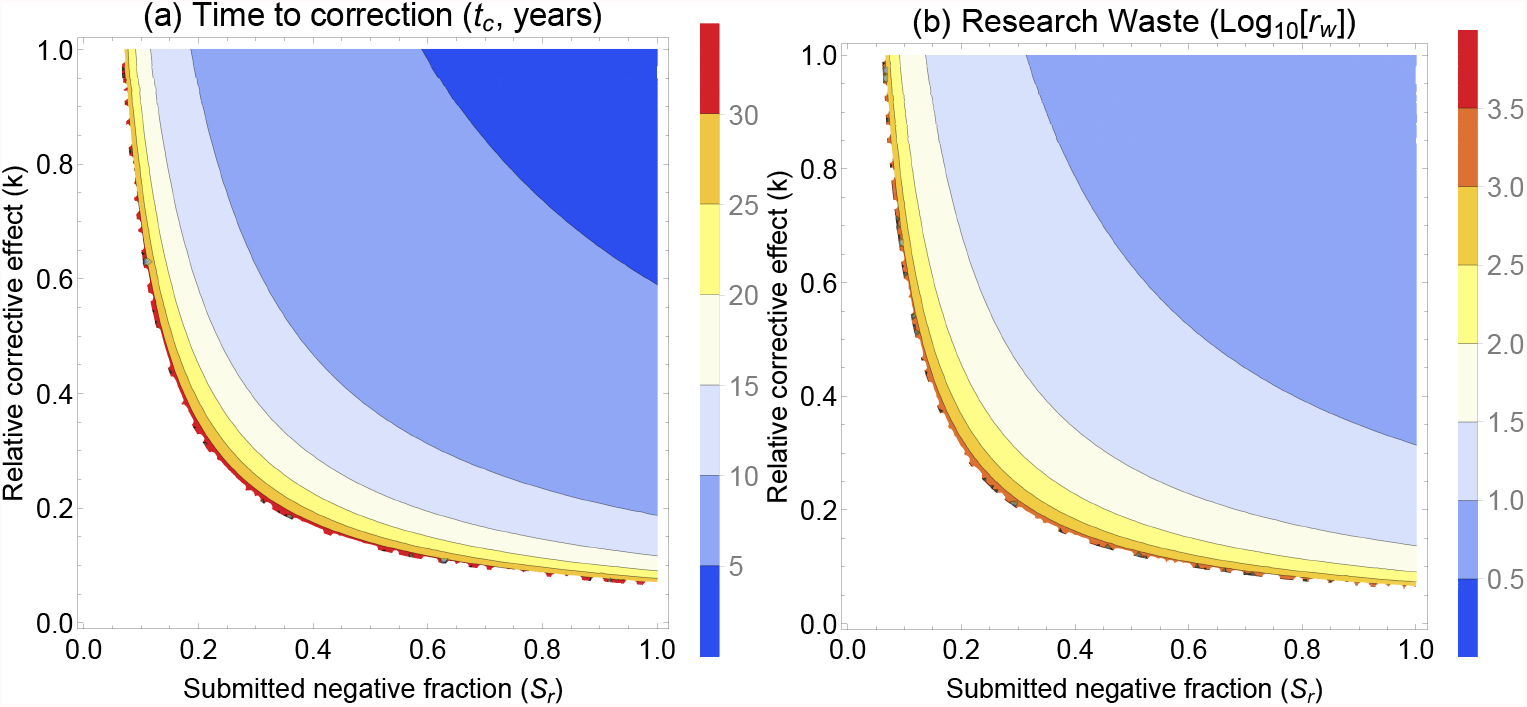
For a spurious result in a field with *f*_*p*_ = 0.10 and *t*_*d*_ = 1 year, (a) Time to correction for a spurious result with varying values of *s*_*r*_ and *k* with all other parameters kept constant (b) The log 10 of research waste (*r*_*w*_) with varying values of *s*_*r*_ and *k* with all other parameters kept constant. Note the log scale on this figure, so that the contour line of 2 is equivalent to *r*_*w*_ = 100, 3 corresponds to *r*_*w*_ = 1000 etc. White areas in the depict regions where no corrective effect is possible by publication alone.

### Hypothetical impacts of eliminating publication bias on research waste and result trustworthiness

With the hypothetical removal of publication bias, figure 5 depicts the likely impacts at varying levels of false positive results. It is immediately apparent that relative to the current paradigm as illustrated previously, correction time, research waste, and corrective effort are all vastly reduced. It can be readily known that were publication bias eliminated, the solution for 4*de* −*g*^2^ ≥ 0 yields 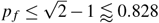. Thus in a situation without publication bias, self-correcting through publication alne could be possible even with up to a 82% false positive rate. This is in stark contrast to the current paradigm.

**Figure 5.**
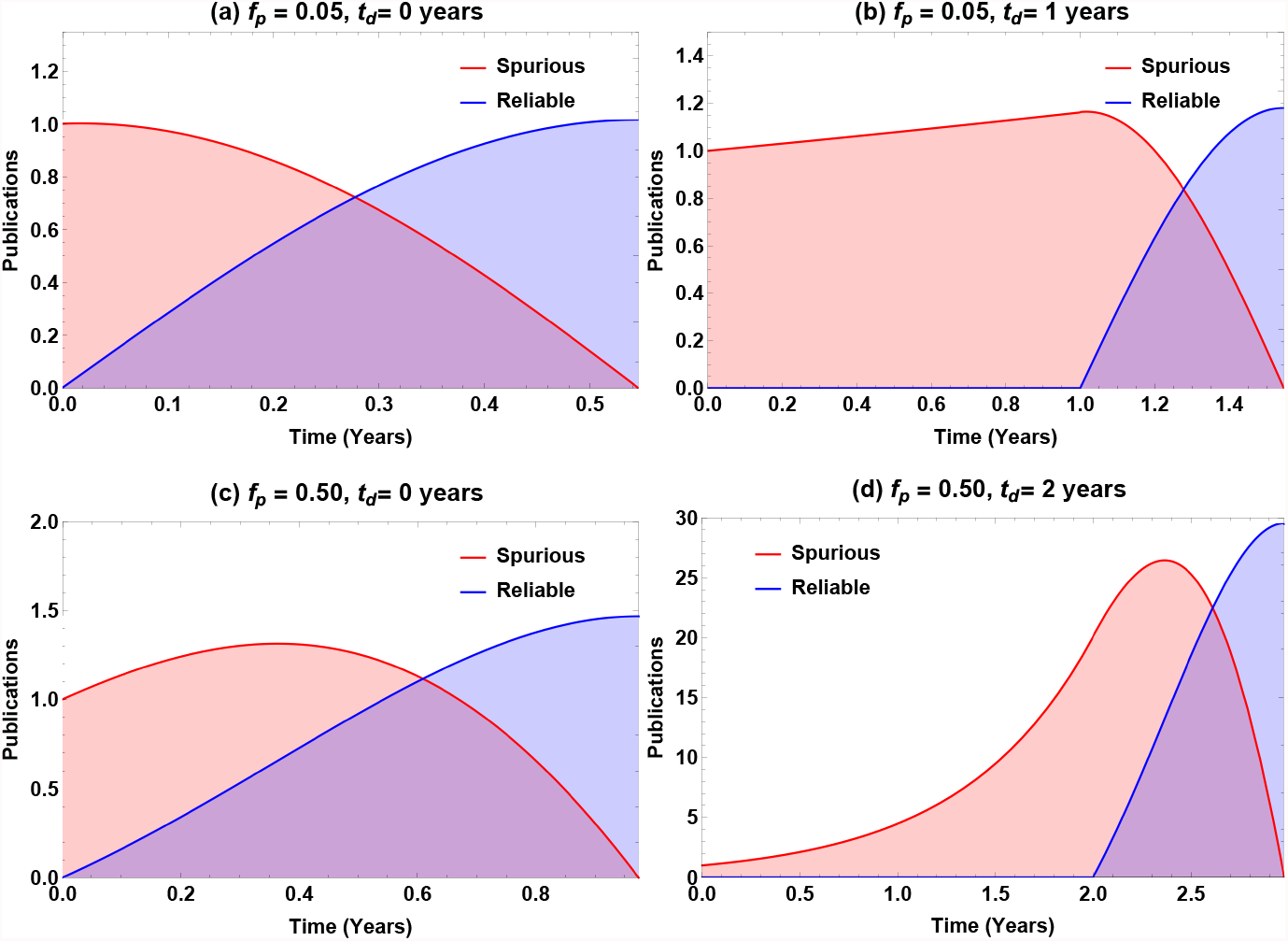
Dynamics of publishing ecosystem in the abscence of publication bias for (a) low false positive rate, no corrective delay (b) low false positive rate, minor corrective delay, (c) very high false positive rate, no corrective delay (d) very high false positive rate, 2 year corrective delay. Note the varying y-axis scales in all subfigures.

## Discussion

The model outlined in this work is a simplification of an inherently complex system, but gives us some predictive insight provided certain reasonable assumptions hold. With current publication incentives, the modelling in this work strongly suggests there are situations where self-correction will be rendered impossible through publication alone. In fields where false positive rates are high, the likelihood of this outcome is increased, and exacerbated by several factors. The time taken for corrective work to begin is a perhaps surprising factor suggested by this investigation. While this might only marginally increase time taken to correction, it has out-sized influence on the volume of research waste produced, and the sheer corrective effort required. This model suggests accordingly that the longer spurious results are ignored, the more subsequent research waste they produce.

The practice of research itself can mitigate this somewhat. Better research conduct and proper statistical analysis yields fewer spurious findings in the first instance. This aside, model results suggest that the file-drawer problem is a substantial one, and that researchers burying their null results is harmful to scientific enterprise. This goes too for the concealing of contradictory results or dis-confirmatory findings. It is critical for healthy science that scientists submit null results, regardless of the perceived status of doing so. Equally, null results should be weighed on par with positive findings. While an exciting result might suggest a research area to pursue, researchers must be mindful to also search for counter-evidence. This is currently not the case, evidenced by the sheer number of citations that retracted publications continue to accrue.

But much of this blame for this be shared with scientific publishing. The fixation on novel, positive results over diligent science creates a perverse incentive to favour even illusory positives over reliable nulls. Fixation on novelty alone is antithetical to healthy science, and journals as the gatekeepers of scientific publication need to acknowledge this and take corrective action of they are serious about upholding scientific trustworthiness. The removal of publication bias has the most stark effect in the model, radically reducing research waste and increasing trustworthiness of findings. This is within the power of journals to implement. Science is from a philosophical standpoint self-correcting, and if journals continue to impede this, they will ultimately be rendered irrelevant.

It is important to note the model outlined here has significant limitations. It is an approximation of complex dynamics, and does not account for factors that might be introduced outside of that simulated ecosystem. In many respects, the model have been overly optimistic about our ability to self-correct. Scientists are humans, subject to human biases. Confirmation bias is inevitable, and volume of evidence may not be enough to dislodge falsified ideas. There is considerable evidence that even if published, more negative findings do not obtain a high level of citation^46^. This is partially accounted for in the parameter *k* in the model, attesting to the relative corrective effect of null-findings, but even this might be overly optimistic. This can lead to a scenario where zombie findings still persist, despite refutation. This might mean that while technical self-correction is achieved, these zombie studies linger on, still creating future research waste. Overcoming this may require a fundamental retraining of researchers^47^.

Another potential limitation of the current model, related to the last point, is that it does not yet consider the potential for “walled-gardens” inside a sub-field. For example, a concept may be refuted by high quality studies, but the falsified idea may persist in a subdomain by citations to the debunked original. This has been seen empirically in certain fields for the trajectories of thoroughly refuted papers^48^, and if common would impede corrective action further. How widespread this is remains unclear, but its net effect in the parlance of this model would be to reduce *k* even further, potentially to the extent that publication-driven self-correction of science is rendered impossible. A recent analysis by Sigurdson *et al*^49^ looks at the specific example of homeopathy, quantifying how the majority of homeopathy trials report positive outcomes for the treatment, despite its central tenets being physically impossible^50^. Over half these studies contained demonstrable statistical errors, suggesting for fields such as this, wall-gardens might mean that no amount of subsequent investigation can dislodge intrinsic pathological science.

There is also a further issue in corrective science - it is incredibly difficult to motivate researchers to undertake such efforts. It has recently been suggested that researchers should get credit for their efforts to correct the literature^51^, and this seems overdue. From personal experience, however, while it is possible to get corrections published in literature^52,53^, it is a common experience to find journals reluctant to engage with good-faith criticism, or even outright hostile to it. Correcting spurious work is a serious effort, and one that goes unrecognised and unrewarded in our current schema. This needs to change if we are serious about scientific quality and research integrity. The model outlined is a useful insight into how self-correction may or may not be achievable, and the factors shaping its efficacy. It also yields insights into how we might mitigate research waste and shape sustainable research in biomedicine. This remains however an understudied problem, and one that demands much more research effort if we are to maintain trust in science, for both the scientific community and the wider public.

## Acknowledgements

DRG would like to thank Profs John Ioannidis, Dorothy Bishop, and Paul Glasziou for their insightful comments and encouragement. He also thanks the Wellcome Trust for their continued support.

## Author contributions statement

DRG conceived the concept, undertook the analysis, and authored the manuscript.

